# Aberrant sorting of hippocampal complex pyramidal cells in Type I Lissencephaly alters topological innervation

**DOI:** 10.1101/2020.02.05.935775

**Authors:** James A. D’Amour, Tyler G. Ekins, Stuti Ghanatra, Xiaoqing Yuan, Chris J. McBain

**Author notes:** Correspondence to: Chris J. McBain (CJM). Author contributions: J.A.D. and C.J.M. designed and conceptualized research; J.A.D., T.G.E., S.G. and X.Y. performed the research; J.A.D. and T.G.E. analyzed the data; J.A.D. and C.J.M. wrote and edited the paper.

## Abstract

Layering has been a long-appreciated feature of higher order mammalian brain structures but the extent to which it plays an instructive role in synaptic specification remains unknown. Here we examine the formation of synaptic circuitry under cellular heterotopia in hippocampal CA1, using a mouse model of the human neurodevelopmental disorder Type I Lissencephaly. We identify calbindin-expressing principal cells which are mispositioned under cellular heterotopia. Ectopic calbindin-expressing principal cells develop relatively normal morphological features and stunted intrinsic physiological features. Regarding network development, a connectivity preference for cholecystokinin-expressing interneurons to target calbindin expressing principal cells is diminished. Moreover, in vitro gamma oscillatory activity is less synchronous across heterotopic bands and mutants are less responsive to pharmacological antagonism of cholecystokinin-containing interneurons. This study will aid not only in our understanding of how cellular networks form but highlight vulnerable cellular circuit motifs that might be generalized across disease states.

## Introduction

Cellular heterotopias within brain structures can result from a variety of developmental insults to an organism and represent breaks from the normal laminar appearance of higher order mammalian brain structures [1, 2]. While heterotopias may arise from diverse causes, they share some common phenotypes [3], offering a unique window of study into the development of cellular networks without the positional cue of layer. Particularly devastating heterotopias involve mutations to genes that encode proteins essential to cellular migration and proliferation [4]. Brains from these patients often appear smooth, lacking the infoldings and gyri of healthy human subjects. Broadly, this condition is referred to as lissencephaly, meaning “smooth brain”. One of the most common and first identified genetic causes of Type I lissencephaly is due to mutations in the Lis1 gene (PAFAH1B1), which encodes an enzyme essential for nuclear kinesis and microtubule stabilization [4, 5, 6, 7]. Unsurprisingly, mutations to other parts of this migratory pathway also result in lissencephalies and recently infections during embryonic development have received renewed attention for their role in microcephalies, such as mosquito transmitted ZIKA virus (for example DCX, 14-3-3 epsilon, RELN, ARX) [3, 8]. These disorders also produce intra-structure cellular heterotopias which are characterized by mispositioned cell somas and disorganized cellular layering. Clearly, mis-lamination is a shared feature of several human neurodevelopmental disorders that merits deeper investigation.

Although rodent brains lack gyri, mice heterozygous for the human mutant Lis1 allele display severe cellular heterotopias in both cortex and hippocampus, developmental defects, hydrocephaly, and enlarged ventricles. These mice also have increased network excitability, lowered seizure threshold, and increased spontaneous mortality rate – features shared with the human condition [9, 10]. In the hippocampus of Lis1 mutants the PCL is often fragmented lengthwise, resulting in multiple PCLs on the deep-superficial axis of the structure, with inter-PCL spaces between. In these mutants, the PCL splits into two distinct bands of excitatory principal cells, a deep and superficial cell layer, although this splitting can be variable and often scattered in appearance.

In light of recent studies suggesting specified microcircuitry among deep versus superficial principal cells and local basket cells in wild type CA1, we wondered if the two heterotopic cell layers observed in Lis1 mutants reflected a functional distinction between discrete microcircuitry of the PCL [11, 12, 13, 14]. Recent evidence suggesting a preferential connectivity between principal cells and either parvalbumin (PV) or cholecystokinin (CCK) expressing interneurons, depending on the extrahippocampal projection target, somatic position of the principal cell, or marker expression of the principal cell, suggests an underlying blueprint in the establishment of hippocampal circuitry and connectivity that has been previously underappreciated in what otherwise appears as a monolithic excitatory lamina, the principal cell layer (PCL) [15, 16, 11, 12, 13, 14, 17, 18]. To what extent are innate wiring motifs disrupted under heterotopia?

Remarkably, in subjects suffering from cellular heterotopias that survive into adulthood cellular networks function surprisingly well and animals are often behaviorally indistinguishable from normal type littermates [19, 20, 21]. In the more thoroughly studied Reeler mouse model, that displays severe cortical and hippocampal mis-lamination, cells in cortex appear to be relatively healthy and are integrated into the local network [22, 23, 24, 25, 20]. Collectively the evidence suggests that the formation of functional synaptic connectivity has some innate resilience to mis-lamination and layers may play little to no role in the guidance and establishment of synaptic connectivity [26, 27, 24, 21]. Furthermore, if there was logic behind the dividing of these heterotopic cell populations in the Lis1^+/−^ mouse it would represent an ideal model to assay the resilience of genetic network formation blueprints to the developmental/local-environment cues of intra-structure position and layering [28, 29, 30]. This might permit us to determine over what relative distances genetic wiring programs are able to locate and synapse on the appropriate postsynaptic targets and shed light on what appears to be intertwined parallel circuitry for information processing in CA1, or identify synaptic connectivity motifs that are more susceptible to heterotopia than others [31, 17]. Ultimately these studies provide key insight into what exactly is the role of layers in higher mammalian brain structures and highlight the proper areas of study for future treatment of cellular heterotopias.

## Methods

### Animal care and breeding

All experiments were conducted in accordance with animal protocols approved by the National Institutes of Health. Lis1^+/ fl+^ male mice (provided by the laboratory of Anthony Wynshaw-Boris, Case Western Reserve University) were crossed with Sox2-cre females (provided by National Human Genome Research Institute transgenic core, Tg(Sox2-Cre)1Amc/J). Sox2-cre females display cre-recombinase activity in gamete tissues, allowing us to genotype and select non-conditional Lis1^+/−^ mutants without the cre allele in one cross. These mice were bred to wild-type C57BL/6J mice (Jackson Labs stock no. 000664) and used for experiments. Both male and female Lis1^+/−^ mice were used for recording and immunohistochemical experiments. Female Ngn2-Cre:RCE (provided by the laboratory of Rosa Cossart, INSERM Marseille, France) mice were used for cell birth-dating experiments. Calbindin-cre mice were obtained from Jackson laboratories (stock no. 028532) and bred to Ai14 animals from also from Jackson (stock no. 007914).

### Cellular birth-dating

Timed pregnancies were established between Lis1^+/−^ males and tamoxifen inducible Ngn2-CreER^TM^:RCE females. Tamoxifen administration in these pregnant mice induces cre-recombination and subsequent eGFP expression in newly born neurons of developing mouse pups. Pregnant mothers were gavaged with tamoxifen (Sigma no. T5648) in corn oil (200-250 μL, 20 mg/mL) at various embryonic time points spanning days E12-17. Pups were genotyped and grown to P27-32 before perfusion and brain fixation in 4% paraformaldehyde in 0.1 M phosphate buffer for 2-4 hours at room temperature or 12 hours at 4°C. Brains were washed, transferred to 30% sucrose in 1x phosphate buffered saline and stored at 4°C. Sections (50-100 μm) were cut on a frozen microtome and stained for calbindin protein (described below). Coronal hippocampal sections were confocally imaged under 20x magnification on a Zeiss confocal microscope, tiled, stitched in the Zen Black software package and post-hoc analyzed for colocalization of calbindin staining and eGFP expression using the Imaris analysis package (Imaris 9.3.1, Bitplane).

### Immunohistochemistry

Standard staining procedures were used for most of the experiments and have been described previously [32] but briefly, deeply anesthetized mice were transcardially perfused with 50 mL of 4% paraformaldehyde (PFA) in 0.1 M phosphate buffer (pH 7.6). Brains were post-fixed overnight at 4°C, then cryopreserved in 30% sucrose solution. Coronal sections were cut (50 μm) on a frozen microtome. Prior to staining sections are washed in phosphate buffered saline (PBS), blocked and permeabilized with 0.5% triton X-100, 10 % goat serum in PBS for two hours at room temperature while shaking. Primary antibodies are applied overnight at 4°C shaking at the appropriate dilution with PBS containing 1% goat serum and 0.5% triton X-100. The following day sections are washed, and a secondary antibody is applied for one hour at room temperature while shaking at a dilution of 1:1000. For most experiments, a final DAPI staining was also used to show lamina of the hippocampus. Sections are then mounted and cover slipped with Mowiol. Primary antibodies: Calbindin (Millipore polyclonal rabbit, stock no. AB1778, 1:1000; or Swant monoclonal mouse 1:1000, stock no. 300); CCK (Frontier Institutes rabbit, stock no. CCK-pro-Rb-Af350, 1:1000). For quantification of inhibitory puncta the procedure was similar with a few adjustments. Coronal sections (50 μm) of dorsal hippocampus were cut, blocked with 10% donkey serum in 0.5% Triton X at room temperature for 2-4 hours. Primary antibodies were applied in phosphate buffered saline with 1% donkey serum and 0.05% triton X-100 at 4°C for 48 hours. Secondary antibodies were left at room temperature for 1-2 hours, before washing and mounting. Primary antibodies: Gephyrin-mouse (Synaptic Systems, CAT no. 147021, 1:1000), Wolfram syndrome 1 (Wfs1)-rabbit (Protein Tech, CAT no. 1558-1-AP, 1:5000), cannabinoid1-receptor (CB1-R)-guinea pig (Frontier Institutes, CAT no. CB1-GF-Af530, 1:5000), parvalbumin (PV)-goat (Swant, CAT no. PVG 214, 1:5000). Calbindin was visualized by using pups from crosses between Lis1 mutants and Calbindin-cre:Ai14. Anti-donkey secondaries: Jackson Immuno Reseach laboratories Inc., AF 405 mouse (715-476-150), AF 488 rabbit (711-545-152), and AF 633 (706-605-148) guinea pig or goat (705-605-147) for visualization of CB1-R- and PV-positive baskets respectively (all 1:500). Images were captured on a Zeiss 880 confocal under 63x magnification using Zen Airyscan image processing. Between 25-30 Z-axis images were collected at Z-steps of 0.159 μm. Analysis was performed on a Max-IP from the first seven of these steps, accounting for 1.1 μm of tissue thereby minimizing Z-axis problems.

Images were quantified in Imaris 9.3.1 software. Twelve principal cells were selected using the Wfs1 staining – half of which were calbindin positive, and cell somas were traced. Gephyrin puncta (with an approximated size of ∼0.25 μm) were automatically detected in the image and excluded if not within 1 μm of a cell soma. In parallel, inhibitory boutons were automatically detected from a pre-synaptic basket cell marker (parvalbumin in one set of experiments, CB1-R in the other). Inhibitory puncta were filtered for proximity to the post-synaptic gephyrin puncta (1 μm or less), and further filtered by proximity to a principal cell soma (0.2 μm or less). Remaining inhibitory puncta were counted on the somas of six calbindin positive, and six calbindin negative principal cells. Dividing puncta counts on calbindin cells by those on calbindin-negative cells yielded synaptic innervation bias measurements such that counts from 12 cells are used to generate a single data point. – A value less than one signifies an avoidance of calbindin positive targets and numbers greater than one signifies a preference for calbindin positive targets.

### Principal cell reconstructions

Slices with biocytin filled cells were fixed (4% PFA and stored at 4°C) and processed for visualization using avidin conjugated dye. Slices were resectioned (50-100 μm) and DAPI stained so cells could be visualized, and their somatic depth could be assessed within the larger hippocampal structure. After staining, slices were imaged, and files were imported to Neurolucida (MBF Bioscience) cell tracing software. Once traced, data sheets were exported for apical dendrite shapes and connectivity profiles for each cell and processed in a custom python script to generate the LRI and ORI measurements later used for morphological clustering.

### Slice preparation

Young adult mice (P20-40) were anesthetized with isoflurane before decapitation. Brains were immediately dissected in dishes of ice-cold dissection ACSF (in mM): 1 CaCl_2_, 5 MgCl_2_, 10 glucose, 1.25 NaH_2_PO_4_ * H_2_0, 24 NaHCO_3_, 3.5 KCl, 130 NaCl. ACSF was oxygenated thoroughly for 20mins by bubbling vigorously with 95% O_2_ and 5% CO_2_ beforehand. For measurement of cell intrinsic properties whole-cell recordings, mono-synaptic inhibition, and disynaptic inhibition experiments coronal slices were cut (350 μm) using a VT 1200S vibratome from Leica Microsystems. Slices were allowed to recover in an incubation chamber at 35°C in the same solution for 30 minutes. For oscillation experiments, the same extracellular slicing and recording solutions were used, and pipettes contained extracellular solution. Slices were cut horizontally (450 μm) from more ventral hippocampus, as oscillations were often extremely weak or all together lacking from coronal sections. We verified that similar migratory problems with the late-born calbindin population occurred in ventral hippocampus (Figure 7B). Oscillation experiment slices recovered for 15minutes at 35°C before being transferred to a custom interface incubation chamber.

### Whole-cell physiology

For electrophysiological recordings slices were transferred to an upright Olympus microscope (BX51WI) with a heated chamber (32°C, Warner Inst.) and custom pressurized perfusion system (∼2.5 mL/min). Recording ACSF contained the following (in mM): 2.5 CaCl_2_, 1.5 MgCl_2_, 10 glucose, 1.25 NaH_2_PO_4_ * H_2_0, 24 NaHCO_3_, 3.5 KCl, 130 NaCl. Electrodes of 4-6 MOhm resistance (borosilicate glass, World Precision Instruments, no. TW150F-3) were prepared on Narishige (PP-830) vertical pipette pullers. Recording were collected using a Multiclamp 700B amplifier (Molecular Devices) with a Bessel filter at 3kHz and Digitized at 20kHz using a Digidata 1440A (Molecular Devices). Protocols were designed, executed and analyzed using the pClamp 10.4 software package (Molecular Devices). Liquid junction potentials were not corrected for and series resistance compensation was not applied. Series resistance was monitored throughout experiments using a −5mV pulse at the start of each sweep and ranged from 12-32MOhms. Cells were biased to −70mV in current clamp mode, and held at −70, −30, and +10mV in voltage clamp mode depending on the requirements of the experiment. For basic properties and morphological recoveries, electrodes were filled with the following, in (mM): 130 K-glu, 0.6 EGTA, 10 HEPES, 2 MgATP, 0.3 NaGTP, 10 KCl. For monosynaptic inhibition experiments, eIPSCs were recorded at −70 mV using electrodes were filled with (in mM): 100 K-glu, 45 KCl, 3 MgCl, 2 Na_2_ATP, 0.3 NaGTP, 10 HEPES, 0.6 EGTA; yielding an E_cl_ of −27 mV. eIPSCs were evoked by local stimulation for 5-10 minutes until a stable baseline was established, then omega-conotoxin GVIA (1 μM) was applied while eIPSCs were monitored for changes in amplitude. Similar experiments were performed washing in omega-agatoxin IVA (250 nM), with QX-314 (2 mM) added to the internal solution. For feedforward I/E experiments electrodes contained (in mM): 135 Cs-MethaneSO_4_, 5 NaCl, 4 MgATP, 0.3 NaATP, 10 HEPES, 0.6 EGTA, 5 QX-314 chloride salt, giving an E_cl_ of −69.7 mV. Internal solutions were adjusted for a pH of 7.4 using KOH and an osmolarity of 290 mOsm. Biocytin (2mg/1mL) was added to thawed aliquots before use. For feedforward inhibition experiments, pilot experiments where stimulation was delivered in CA3 did not include a wash-in of excitatory blockers as activation of direct monosynaptic inhibition was less likely. For most of the experiments however, stimulation was delivered in the s. radiatum of CA1 and APV (50 uM) / DNQX (20 uM) was added to block glutamatergic transmission, permitting us to determine and subsequently subtract the monosynaptic component of the inhibitory response. These data were pooled. Recordings where IPSCs were not reduced by at least 30% were excluded.

### Extracellular field potentials

For LFP recordings, slices were transferred onto an interface chamber with two manipulator-controlled electrodes positioned under 25x visual guidance. Carbachol (20 μM) was applied to induce slice oscillations. Recordings were made at 10kHz, low and high pass filtered (8 and 100 Hz, respectively) and mean subtracted. Cross correlation was the max real value resulting from the inverse fast-fourier transformation of F_1_ and F_2_; where F_1_ = fft(signal sample from channel 1), and F_2_ is likewise for channel 2, after a flip operation. Cross correlation summary values are the max cross-correlation value in the resulting vector C. The temporal shift between the two signals is the X-coordinate (in milliseconds), corresponding to this cross-correlation peak. Experiments were processed such that channel-1 and channel-2 always corresponded to the same side of the principal cell layer (deep vs superficial).

### Data analysis

Initial data exploration and analysis was performed in custom Python scripts. For further plotting and statistical analysis Graphpad Prism was used for physiological data. For soma positioning measurements and gephyrin puncta quantification, Microsoft excel sheets were used. K-means clustering was performed in Python using the Scikit learn clustering and decomposition packages. Both clustering routines were supervised (Figure 3 and 4), in that they expected K-means n = 2. For morphological clustering this was to replicate prior work and aid in identification of calbindin positive and negative principal cells. For physiological properties, we wished to ask if the two morphological populations might be reflected in our physiology data.

### Statistics

P values represent Welch’s t-tests for comparisons of two independent samples, unless otherwise noted. Student’s paired t-tests were used for intra-sample (like inhibitory puncta) and pre-post wash comparisons. R values represent Pearson’s cross-correlation unless otherwise noted. Quantification and error bars are standard error of the mean.

## Results

### Heterotopic banding of the principal cell layer in Lis1 mutant mice

Non-conditional heterozygous Lis1^+/−^ mice were generated by breeding a Lis1^+/ fl+^ line to Sox2-cre animals. Lis mutants were often noticeably smaller than litter mates. Some animals displayed severe macroscopic brain abnormalities, including enlarged ventricles, hydrocephaly, intracranial bleeding, and spontaneous death. Lis1^+/−^ mutant mice that survived to 3-5 weeks of age were used for experiments and subsequent breeding. In coronal sections from dorsal hippocampus Lis1 mutants displayed heterotopic banding of the principal cell layer (Figure 1A). Banding varied in severity, cell soma density, and septal-temporal extent. Most animals displayed the strongest banding in area CA1, with fewer mice showing multiple PCLs past region CA2. Region CA3 rarely appeared banded, but instead scattered and uncompacted. Mice occasionally had three distinct layers or clustered islands of cells, but most typically two prominent PCLs could be seen (Figure 1A, *right* vs *left*). Deeper bands were typically situated in what would be s. oriens-alveus in a non-mutant animal. In measuring from the border of the alveus and the cortex radially (toward radiatum, known as the radial axis of CA1), the entirety of normal WT PCLs were located between ∼175 – 300 μm. In Lis1 heterozygous mutants, superficial bands were positioned between ∼ 250 – 320 μm and deeper heterotopic bands (positioned closer to the alveus) were between ∼ 100 – 190 μm. Of the two bands, the superficial tended to be more densely populated and closer to the normal positioning of the PCL in normal type mice (Figure 1 A and B). We next considered whether these heterotopic bands were splitting randomly, or if the banding represented distinct cell populations.

**Fig 1.**
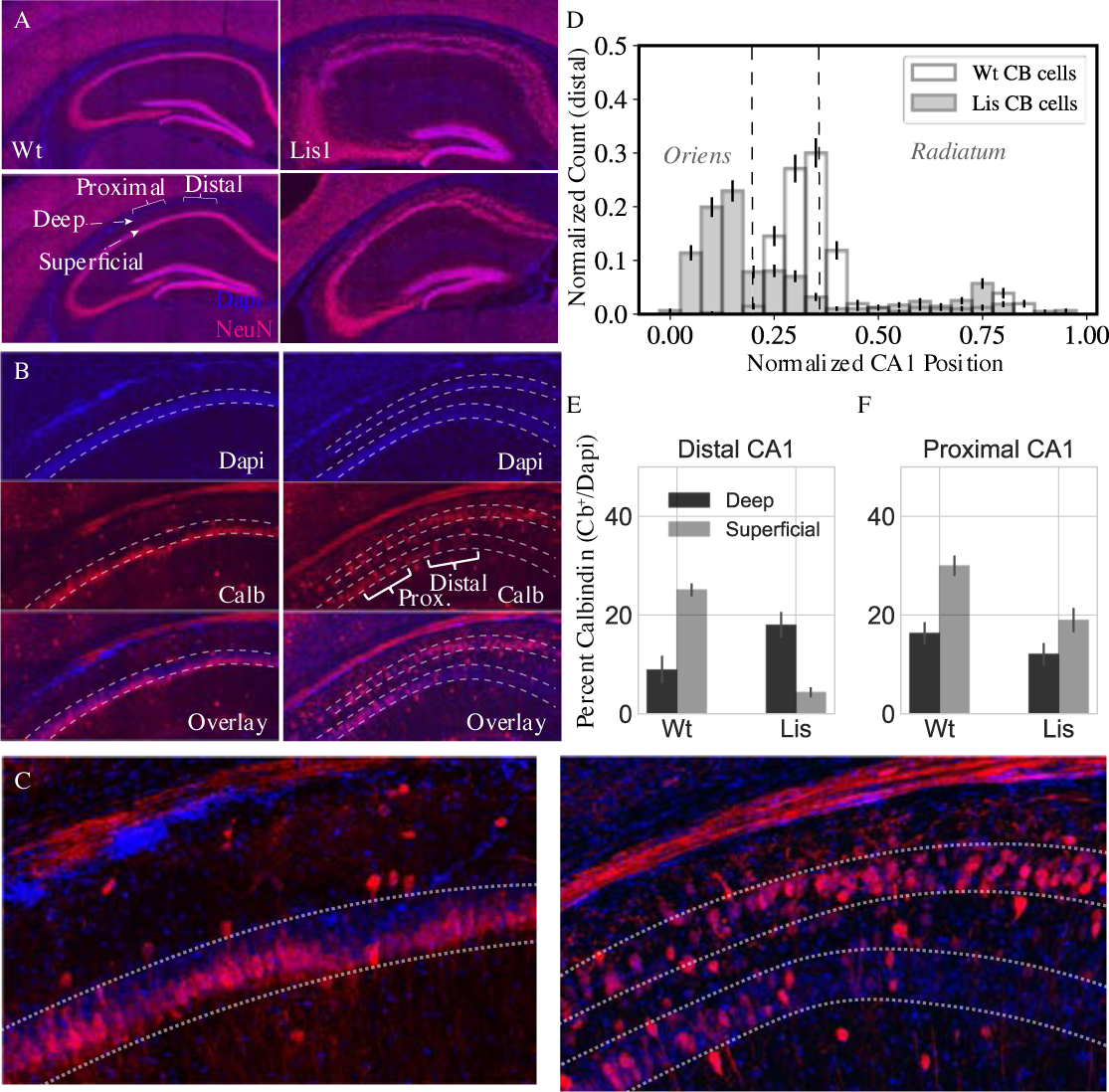
Lis1^+/−^ mice display heterotopic banding and ectopic po<u>s</u>itioning of calbindin-expressing principal cells. (A) *Left*, two coronal NeuN stained images from differing levels of dorsal CA1 hippocampus in a non-mutant littermate. *Right*, approximately matched coronal sections from a Lis1 mutant displaying heterotopic banding of the PCL. (B) *Left*, non-mutant and mutant (*right*), staining of CA1 highlighting the position of the PCL, calbindin-positive neurons, and overlay. Note the deep layer preference of calbindin-expressing neurons, particularly in distal CA1 in mutant. (C) Higher magnification view of the overlay images in (B), for non-mutant (*left*) and mutant (*right*). (D) Normalized histogram showing the positioning of calbindin-expressing cells in mutants with PCL banding compared to non-mutant mice. (E) Percentage of cells in deep and superficial layers expressing calbindin in distal CA1 (for normal mice, the single PCL is divided in half radially). Counts represent number of identified calbindin soma divided by number of DAPI identified cells, Wt: deep 8.9 ± 2.8 %, superficial 25.1 ± 1.3 %; Lis1^+/−^: deep 18.0 ± 2.8 %, superficial 4.4 ± 1.0 %. (F) Same as (E) for proximal CA1, Wt: deep 16.3 ± 2.3 %, superficial 30.0 ± 2.0 %; Lis1^+/−^: deep 12.1 ± 2.3 %, superficial 19.0 ± 2.4 %; n = 12 Wt and 12 Lis1^+/−^ slices for distal and 12 and 11 for proximal, from 6 animals.

### Calbindin expressing principal cells preferentially position in the deeper heterotopic band of CA1 in Lis1 mutants

In order to better understand the banding process in heterozygous Lis1 mutants, immunohistochemistry experiments were carried out for principal cell markers and quantified by normalized expression levels in each heterotopic band (*n* antibody stained cells / *n* dapi cells in same region of interest). In addition to marking a subpopulation of GABAergic cells, Calbindin is expressed in superficial principal cells in several species [13]. Consistent with these reports, our Lis1 normal type litter mates show calbindin-expression among superficial principal cells of CA1 (Figure 1B, *left*). These cells are tightly packed, forming one-three rows of somas on the s. radiatum adjacent (superficial) side of the PCL. Conversely, calbindin staining in Lis1^+/−^ mice showed a strong bias for calbindin-expressing principal cells to occupy the deeper heterotopic principal cell layer (Figure 1B, *right*). Figure 1D shows a normalized histogram of identified calbindin-positive cell soma positions in Lis1 mutants and litter mate controls. For quantification and comparison purposes, the single wild type PCL is divided in half radially and analyzed as separate deep and superficial bands (Figure 1E; Distal CA1 Wt: deep 8.9 ± 2.8 %; superficial 25.1 ± 1.3 %; Lis: deep 18.0 ± 2.8 %; superficial 4.4 ± 1.0 %, n = 12 Wt and 12 Lis1^+/−^ slices respectively from 6 animals). This finding was not a general feature of having the Lis1 mutant allele, as in animals with less severe banding (or in the same slices nearer CA2) but still carrying the mutant Lis allele, the PCL displayed relatively normal, superficial calbindin soma positioning (Figure 1F; Proximal CA1 Wt: deep 16.3 ± 2.3 %; superficial 30.0 ± 2.0 %; Lis: deep 12.1 ± 2.3 %; superficial 19.0 ± 2.4 %, n = 12 and 11, respectively). Given that principal cells are generated near what becomes the alveus and migrate radially during embryonic development in a deep to superficial manner [33, 23, 34], the calbindin staining pattern suggested a late born population undergoing migratory failure in the Lis1^+/−^ mouse.

### Embryonic development of the calbindin expressing principal cells

Superficial principal cells in normal mice arise near the end of gestation (Emb days 16-17) [33, 23, 13]. Our initial data suggests that the heterotopic banding in Lis1^+/−^ mice may arise from a migratory stalling event, where later born superficial-preferring cells were unable to overcome a migratory burden and instead form a new deep heterotopic layer. In order to test this hypothesis and ensure that a novel population of deeply positioned principal cells was not adopting calbindin expression in Lis1^+/−^ mice, cellular birth-dating experiments were performed.

In timed pregnancy experiments using Lis1 mutants crossed to Ngn2-cre:RCE mice, tamoxifen administration induces cre-recombination and subsequent eGFP expression in newly born neurons of developing mouse pups. Pregnant mothers were gavaged at various embryonic time points spanning days E12-17. After pups were born, they were perfused and fixed at ∼P30 for calbindin staining, and subsequent quantification of the percentage of eGFP expressing neurons from any time point that were co-stained for calbindin (Figure 2A-C). Approximately 10% of cells born on E12-E13 expressed calbindin at P30 (Figure 2D; Wt: 9 ± 3 %; Lis: 12 ± 3 %, n = 95 cells and 168 labelled cells analyzed from 5 animals, respectively) in both Lis1 normal type littermates and mutants. Cells born E14-E15 co-stained for calbindin 42 ± 9 % of the time for wild type and 52 ± 8 % (n = 221 and n = 128, from 10 animals) for Lis1 mutants and cells born E16-E17 co-stained for calbindin 54 ± 7 % of the time for wild type and 71 ± 9 % for Lis1 mutants (n = 48 and 20 RCE labelled cells from 11 animals). While the timing of calbindin cell birthdates remained similar to littermate controls in Lis1^+/−^ animals in that calbindin cells arise late in embryonic development (Figure 2D), positioning of these cells differed substantially. Later born cells positioned more superficially in normal type littermates (smaller PCL depth measurements), while they positioned more deeply in mutant mice (Figure 2E and F, E represents counts from single experiments data are averages and summarized in F). These results suggest that deeply positioned calbindin expressing cells in the Lis1 mutant are the same late-born cell population that are now ectopically positioned in the deeper heterotopic band.

**Fig 2.**
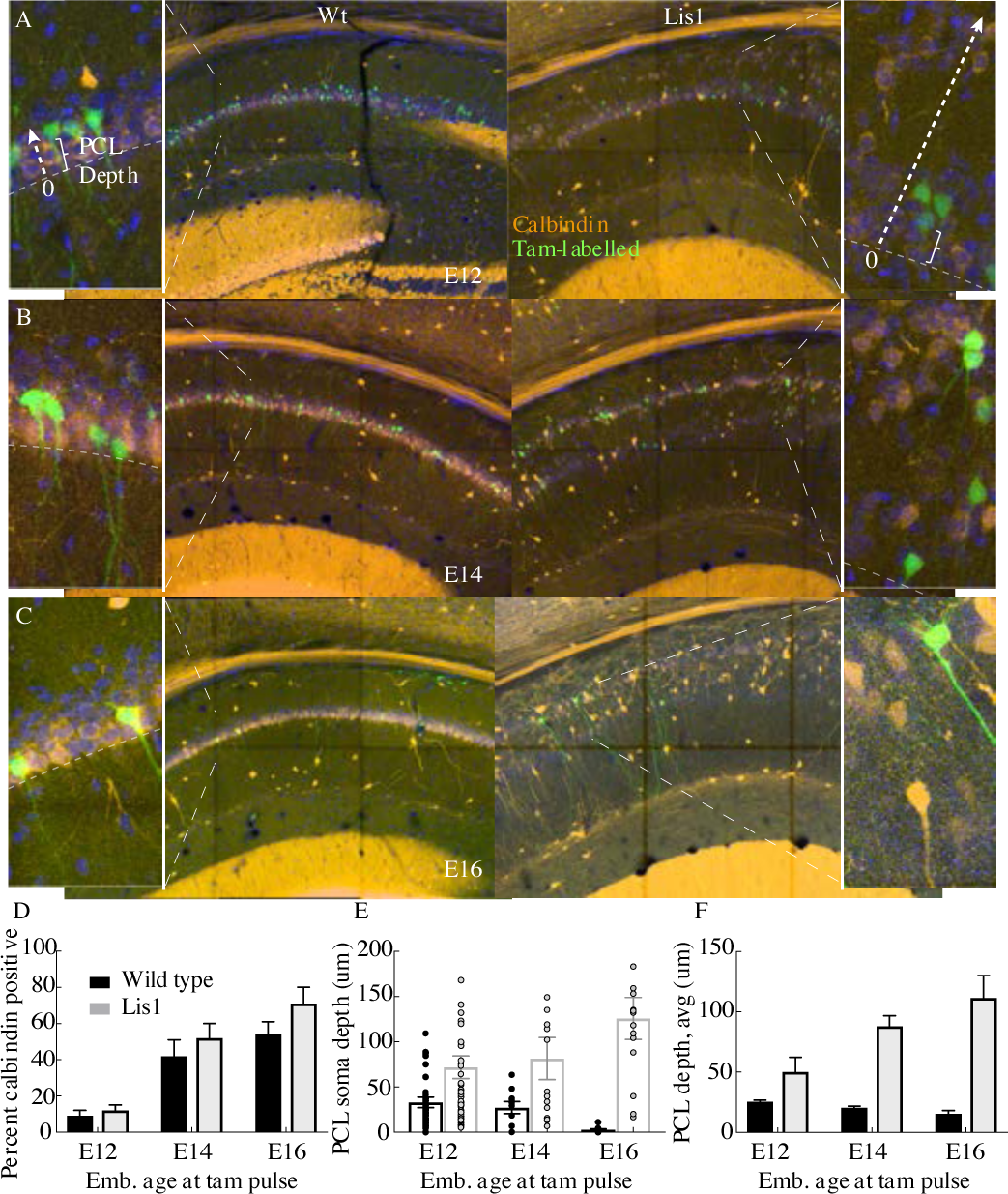
Cellular birth-dating indicates ectopic Lis1^+/−^ calbindin cells are the same late-derived embryologic population. (A) Non-mutant (*left*) and mutant (*right*) example birth-dating images for a litter tamoxifen dosed between E12-E13. Note the cutout, displaying how cellular somatic positioning was measured from the front of the PCL (as opposed to normalized structural position). Green corresponds to cells born during tamoxifen administration; orange is calbindin immunohistochemistry staining carried out when litters are P30. (B) Same as in (A) but for litters dosed at E14-E15. (C) Same as (A) but for litters dosed at E16-E17. (D) Quantification of the fraction tamoxifen-marked neurons co-staining for calbindin antibody from each timepoint. E12: Wt: 9 ± 3 %; Lis1^+/−^: 12 ± 3 %, E14: Wt: 42 ± 9 %; Lis1^+/−^: 52 ± 8 %, E16: Wt: 54 ± 7 %; Lis1^+/−^: 71 ± 9 %. (E) Example counts from single images at each timepoint for PCL depth measurements. Later born cells position more superficially (front of the PCL) in non-mutants, but deeper in Lis1^+/−^ littermates. (F) Group averages for the measurements shown in (E). E12-Wt: 25.4 ± 1.4 μm; Lis1^+/−^: 50 ± 12.3 μm, E14-Wt: 20.5 ± 1.3 μm; Lis1^+/−^: 88 ± 8.7 μm, E16-Wt: 15.6 ± 2.6 μm; Lis1^+/−^: 111.6 ± 18.3 μm.

### Calbindin-expressing principal cells retain a complex apical morphology

Previous studies have documented variation in CA1 principal cell morphology, particularly in comparing basal and apical dendritic trees [35, 11, 36]. These morphological features can be reliably used to differentiate excitatory neuron subtypes. In particular, calbindin-expressing principal cells have more complex apical dendritic trees (more branching), than calbindin-negative counterparts [36]. This has enabled offline characterization of excitatory cell group through K-means clustering of morphological features after cellular reconstructions. A prior study using this approach suggested that clustering was greater than 90% accurate as verified by mRNA and in-situ hybridization approaches but comes with the drawback that every recording must be histologically processed, virtually reconstructed, and analyzed [36]. Additionally, there is a minimal threshold for the amount of dendritic tree that must be recovered and drawn for clustering to be accurate.

In our 63 best recovered principal cells morphologies from physiological recording experiments (n = 32 wild type, n = 31 Lis1^+/−^) we implemented a k-means clustering algorithm based on dendritic branch connectivity and lengths, as done previously [36]. The clustering results from Lis1 mutant and normal type litter mate cellular morphologies are shown in Figure 3A and B. The same process was applied to mutant and normal cells, but these groups were processed independently by a supervised k-means algorithm that expected two groups, corresponding to complex and simple morphologies. The data show that relatively simple and complex cell morphologies persist in the Lis1 mutant, in approximately the same proportions to normal type mice, with nearly overlapping cluster centers (complex cells, Wt: [−0.1, 0.8], Lis1: [−0.4, 0.9]; simple cells, Wt: [−1.8, −1.3], Lis1: [−1.7, −1.2]). A visual comparison of some of the more obviously simple and complex cellular recoveries suggests the sorting has been successful (Figure 3A & C, note deeply positioned complex cells in Lis1 mutant). Grouping cells by the assigned shape cluster and plotting the associated PCL depth measurements (from the border between the first PCL and the radiatum) gives further support to the sorting results, as complex cells were located superficial to simple cells in normal type controls and scattered but generally deeper than simple cells in Lis1 mutants, in agreement with our calbindin staining experiment (Figure 3D PCL depth, Wt complex: 35 ± 8.0 μm, simple: 50.2 ± 6.3 μm, n = 13 and 11; Lis, complex: 127 ± 23.4 μm, simple: 94.6 ±12.3 μm, n = 8 and 13). Additionally, we observed that complex cells in both mutant and non-mutant animals tended to have their first prominent apical branch bifurcation points sooner than simple cells (Figure 3E). This suggests that the complex apical branch morphology can still be used to identify putative calbindin-expressing principal cells in Lis1 mutants. More traditional morphological analyses such as sholl intersections fail to show clear differences between complex and simple cell types when they are pre-sorted by K-means label, highlighting the usefulness of analyzing branching patterns with this approach (Figure 3F-H, note that sholl intersections represent apical dendritic trees only and do not include basal dendrites).

**Fig 3.**
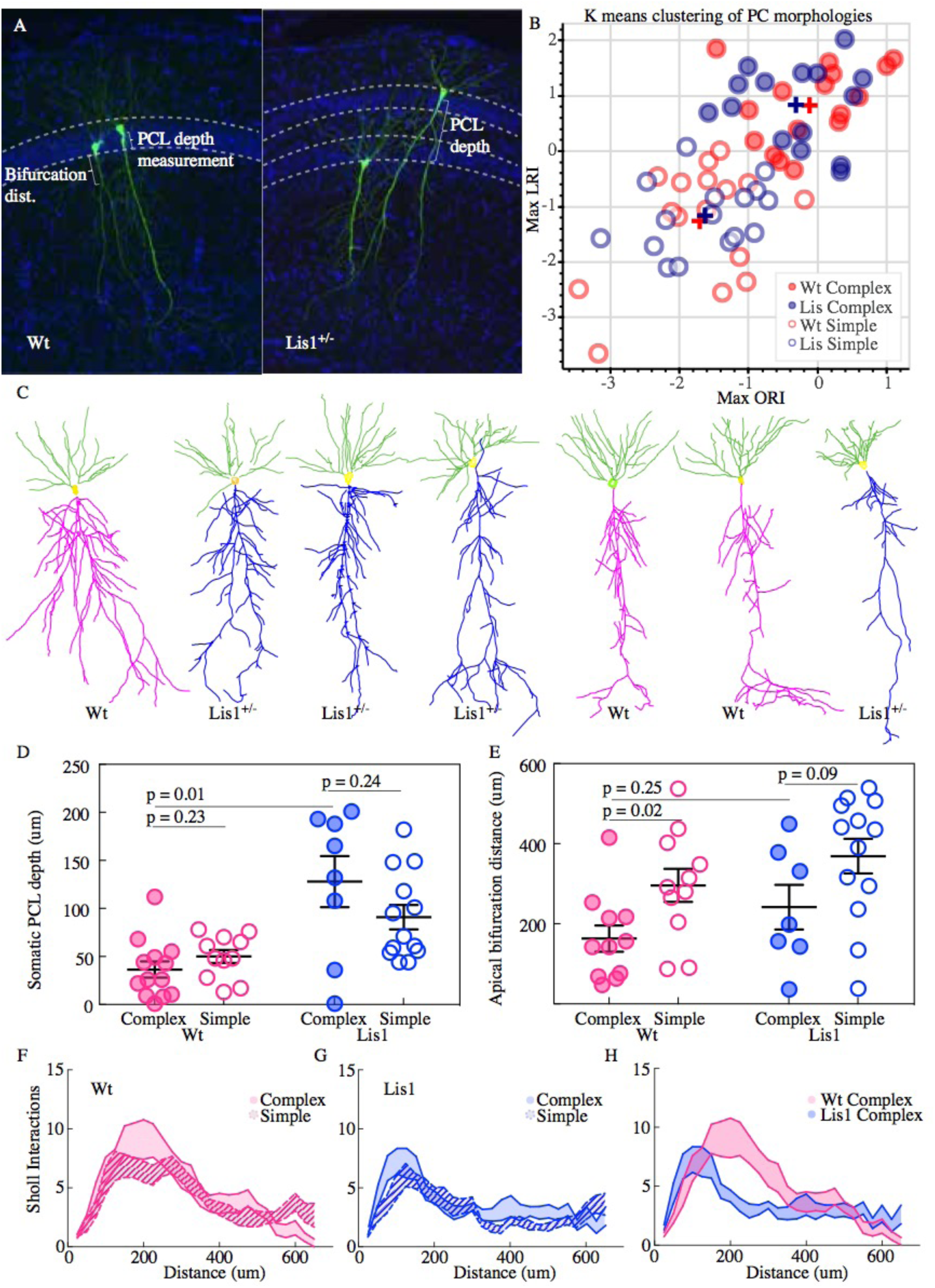
Lis1^+/−^ calbindin-expressing PCs retain relatively complex morphologies. (A) Recovered cells from non-mutant and mutant experiments, highlighting different apical dendritic morphologies, complex and simple. Complex morphologies have been previously shown to be highly predictive of calbindin expression (Yiding et al., 2017). (B) Supervised K-means plots (63 best recovered cells, cluster num. = 2) carried out separately for mutant and non-mutant data (blue and red respectively). Filled circles correspond to complex morphologies and open circles are simple. (C) Example morphological reconstructions, ranging from most complex (*left*) to simple (*right*). (D) Positional properties for predicted calbindin (complex, filled circles) and non-calbindin expressing (simple, open circles) principal cells. Note predicted calbindin expressing cells were superficial to non-calbindin predicted, and this trend was inverted for Lis1^+/−^ mutants. Wt: complex 36.42 ± 8.5 μm, simple 50 ± 6.9 μm; Lis1^+/−^: complex 128 ± 26.6 μm, simple 90.9 ± 12.7 μm, n = 13, 11, 8, 13, respectively. Depth is measured as it was for Fig 2 from the front/superficial side of the PCL. (E) Group sorted measurements for distance along primary apical dendrite until first prominent bifurcation occurs. Wt: complex 163 ± 32.8 μm, simple 295.9 ± 41.4 μm; Lis1^+/−^: complex 241.6 ± 55.8 μm, simple 368.9 ± 43.4 μm. Note complex cells tend to bifurcate sooner in both mutant and non-mutants, though some Lis1^+/−^ complex cells begin to show longer bifurcation measurements. (F) Sholl interactions from Wt apical dendrites alone, of complex and simple sorted cells. (G) Likewise, for Lis1 mutants. (H) Overlay of the complex morphology sholl data from non-mutant and mutant experiments. Despite retaining a relatively complex population, complex Lis1^+/−^ principal cells have decreased apical dendritic branching that peaks closer to the soma.

Importantly, sholl analysis of complex cells from Lis1 mutants and complex cells from control litter mates revealed a reduction in branch intersections in Lis1 complex cells (Figure 3H). While non-mutant complex cells typically have peak sholl intersections of 8-11 around 200 μm from the soma, Lis1 mutant complex cells have fewer peak intersections (∼7), closer to the soma (∼125μm) (Wt n = 10 complex and 14 simple; Lis n = 10 complex and 12 simple). While relatively speaking, the complex and simple subtypes persist in the Lis1 mutant, there has been an effect of the mutation, either direct or indirect, in stunting general morphological development.

### Lis 1 mutants display disrupted physiological properties

From the whole-cell recordings that were used for morphological reconstructions in Figure 3 a battery of intrinsic physiological properties were analyzed in two ways. Several of these properties are shown in Figure 4. Each property was plotted against the PCL depth of the soma (somatic depth from the radial side of the PCL) from which the recording was made (Figure 4A). The same data were also sorted into putative calbindin-positive and calbindin-negative cell types as predicted by either complex or simple morphologies (Figure 3B & 4B). Resting membrane potential displayed a pearson r value of 0.44 for correlation with position in normal type litter mates, and a r-value of 0.07 in Lis1 mutant mice (Wt: n = 23, Lis: n = 23). Sag index correlated with position at an r-value of 0.5 in normal mice and an r-value of 0.05 in Lis mutants (Wt: n = 24, Lis: n = 26). Input resistance and depth in normal type mice had a correlation value of r = 0.2, while in Lis mice r = −0.1 (Wt: n = 23, Lis: n = 26).

**Fig 4.**
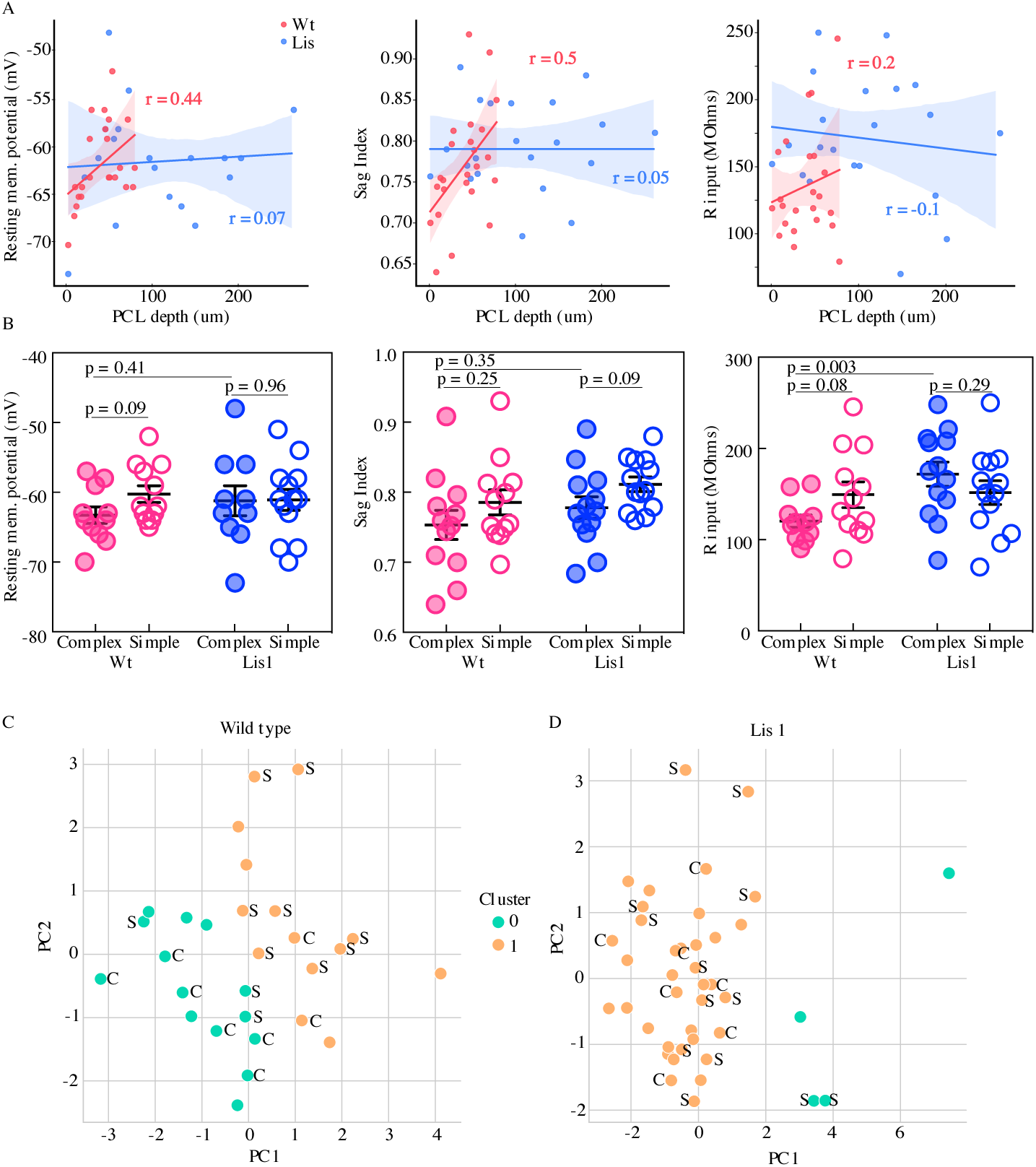
Physiological properties of calbindin positive and negative morphological clusters. (A) *Left*, somatic PCL depth correlations with cellular resting membrane potential for non-mutant (red) and mutant (blue) recordings. *Middle*, likewise, for sag index, where values closer to 1 correspond to less sag exhibited. *Right*, same for input resistance. (B) Same data as in (A), grouped by predicted calbindin expression. (C) Supervised K-means (n = 2) sorting wild types. A handful of electrophysiological properties alone are capable of reasonably accurate morphological subtype prediction (and therefore calbindin expression). C’s and S’s correspond to the data points associated morphological group, note that even mis-categorized points are near the midline. Of 8 morphologically complex cells, 6 are found in in physiological cluster 0, of 11 simple cells, 8 are found in physiological cluster 1. (D) Same as in (C) for Lis1 mutant recordings. Physiological properties are less capable of predicting morphological cluster in Lis1 mutants.

In sorting recorded data by putative cell type, we noted that many of the positional differences observed in Figure 4A persisted or at least trended toward significant in normal type littermates (complex cells are filled circles, open are simple; Resting membrane potential: Wt mean complex −63.3 ± 1.2 mV, simple −60.3 ± 1.2 mV, p=0.09 n = 11 and 12; Sag index: mean complex 0.75 ± 0.02, simple 0.79 ± 0.02, p=0.25 n = 12 and 12; Input resistance: complex 120.4 ± 6.8 MΩ, simple 149.3 ± 14.11 MΩ, p=0.08 n = 11 and 12). Some of these differences in sub-types were still detectable in Lis1 mutants, but differences between sub-types for most properties seemed substantially reduced from normal mice (RMP: mean complex −61.2 ± 2.1 mV, simple −61.1 ± 1.5 mV, p=0.96 n = 10 and 13; Sag index: mean complex 0.78 ± 0.02, simple 0.81 ± 0.01, p=0.09 n = 13 and 13; R input: mean complex 171.9 ± 13.2 MΩ, simple 123.3 ± 13.02 MΩ, p=0.29 n = 13 and 13).

We wondered if there were physiological subtypes of principal cells and how those subtypes might correspond to our previously identified morphological subtypes. Principal component analysis and subsequent K-means clustering was carried out on the physiological data (Figure 4C and D, resting membrane potential, sag index, input resistance, spike amplitude, adaptation ratio, firing frequency at 2x threshold, spike threshold, and after hyperpolarization amplitude were used for physiological clustering). We then scored where morphologically identified cells fell in the physiological clusters. Out of eight morphologically complex cells, six were found in physiological cluster 0 and the remaining two in physiological cluster 1. Of eleven morphologically simple cells, eight were located in physiological cluster 1 and the remaining three in cluster 0, suggesting that these physiological clusters roughly correspond to the two morphological subtypes identified in Figure 3 for normal type littermates (Figure 4C). The same analysis in Lis1 mutants yielded uneven cluster counts, and no clear relationship between physiological cluster and morphological cluster (Figure 4D).

### Basket cell – principal cell innervation biases are differentially affected in the Lis1 mutant hippocampus

Having gained insight into how the heterozygous Lis1 mutation impacts the development of principal cell properties of positioning, embryonic birthdate, morphology and intrinsic physiology, we next wondered how ectopic calbindin cells were integrated into the local synaptic network of CA1. Prior studies have suggested a preferential and complementary innervation bias among two types of local basket cells found in the CA1 subfield – parvalbumin-containing (PV) and a subset of cholecystokinin-containing (CCK) inhibitory interneurons. PV-expressing basket cells preferentially innervate deeply situated calbindin negative principal cells, while CCK-expressing interneurons have a similar bias, but for superficial calbindin positive principal cells [11, 14, 17]. We wondered if these innervation patterns were present in the Lis1 mouse despite ectopic cellular layering, which might shed light on how positioning and layering effect synaptic network development of brain structures.

To begin to assay this network feature in our Lis1 mutants we first asked where these two types of basket interneuron somas were positioning in mutant mice. Immunohistochemical staining experiments were performed using antibodies against PV and CCK (Figure 5A and B). The soma of stained interneuron classes are plotted in binned and normalized histograms in Figure 5B, left and right for PV and CCK, respectively (filled bars for controls dashed bars for mutants). Vertical dotted lines show the approximate location of the wild type principal cell layer. Note that for this figure, somatic position is measured from the alveus/cortical border toward the s. radiatum across the entire radial depth of CA1, as opposed to how it is measured when examining principal cell layer depth (compare with Figure 3F & 4A). As these interneurons often position on the edges of, or outside of the PCL this measure is more appropriate when assessing the overall radial structure of CA1. Our data indicate that both PV- and CCK-containing cell types have undergone superficial radial shifts, that is, the cell bodies have moved towards the s. radiatum. Notably, this is opposite the direction in which calbindin positive principal cells are shifted in Lis1 mutants (Figure 1 & 2). Overall PV-containing somatic shifts appear less severe than CCK-containing shifts, but in both cases a few drastically shifted somas were observed (right tail of dashed histograms).

**Fig 5.**
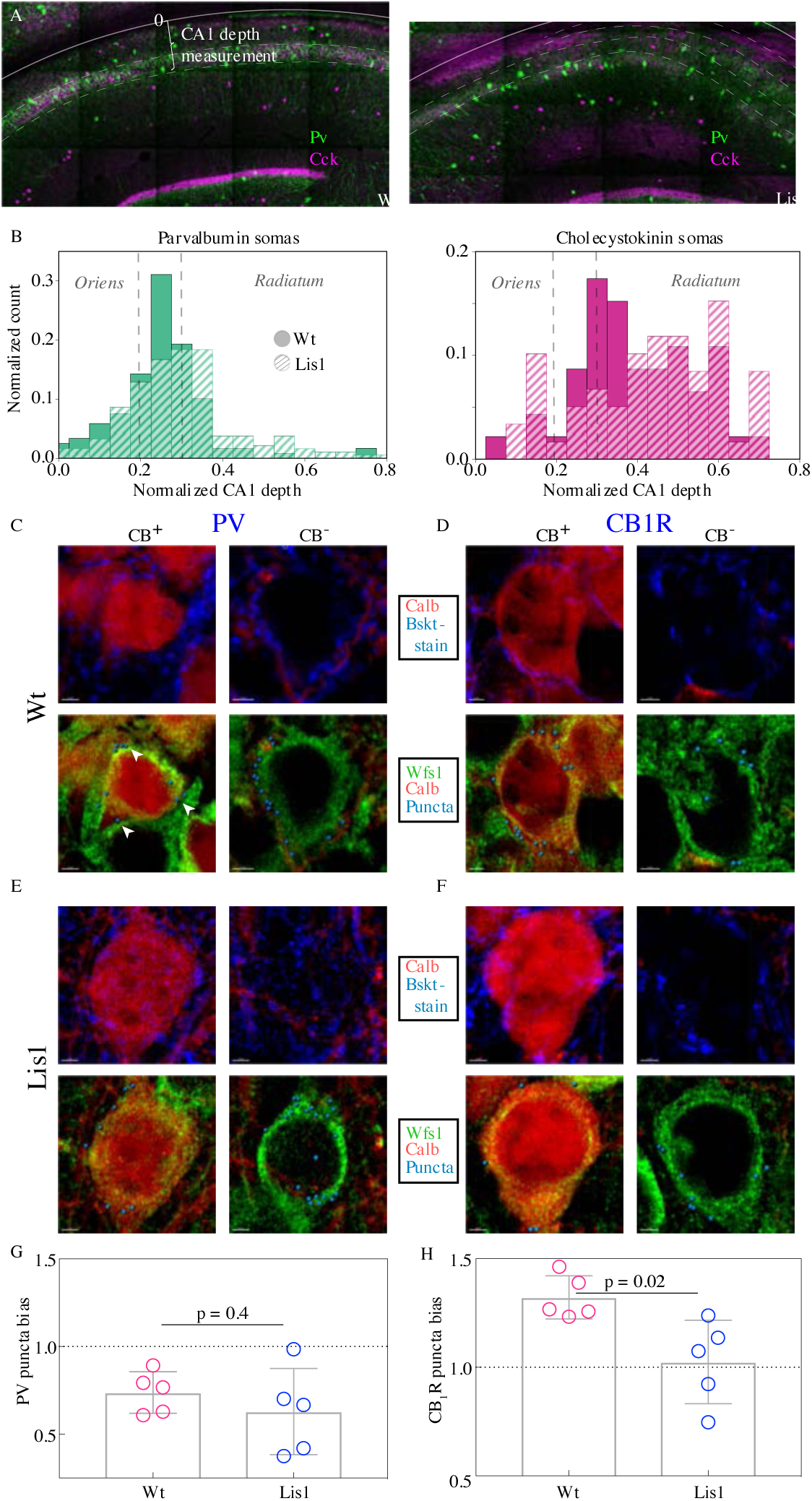
CCK-expressing basket cells have decreased innervation preference with ectopic calbindin positive principal cells. (A) Low magnification images showing the locations of parvalbumin and cholecystokinin-expressing interneurons in the CA1 hippocampus. Note the CA1 depth measurement from the back of the oriens – this measure is more appropriate for assessing somatic position within the larger CA1 structure, as opposed to PCL depth used elsewhere in the study. (B) Normalized histograms of basket cell soma depth measurements along the radial axis of CA1, both PV- (*left*) and CCK-containing (*right*) inhibitory interneuron somas show modest superficial shifts in Lis1^+/−^ mice. (C) High magnification images of a staining experiment for the quantification of PV-containing inhibitory puncta from control littermate samples. *Left*, an example CB-expressing principal cell. *Right*, an example non-CB-expressing principal cell. The top row shows calbindin and parvalbumin staining, the bottom row shows the same cells with calbindin, Wfs1 staining which was used to draw the cell border, and the puncta derived from the parvalbumin staining shown above (arrows point to a few in the first panel) – these puncta are filtered for proximity to a postsynaptic gephyrin puncta (channel not shown). (D) Same as in (C), except the interneuron staining is for the cannabinoid receptor 1, highly expressed in the terminals of CCK-expressing interneurons. (E & F) Same as the corresponding above panels, but for samples from Lis1 mutant littermates. (G) PV puncta bias summary. PV puncta had a modest preference for non-calbindin expressing principal cells in both non-mutant and mutant slices. PV-calbindin preference: 0.74 ± 0.05 and 0.63 ± 0.11 innervation biases for normal type and mutants respectively, p = 0.55, each point represents 12 cells from a slice, n = 3 pairs of littermates from 3 litters. (H) Same as in (E), but for experiments where the PV antibody was replaced by the CB1-R antibody. Non-mutant CCK baskets displayed a preference for calbindin-expressing principal cells that was lost in Lis1^+/−^ mice. CB1-R-calbindin preference: 1.32 ± 0.04, 1.02 ± 0.09 for normal type and mutant respectively, p = 0.02. Scale bars for C-F are 2 μm.

To begin to probe synaptic network development under heterotopia we performed high magnification immunohistological staining experiments with four simultaneously visualized channels (Figure 5C - F). This permitted the identification of inhibitory synapses on the somas of calbindin-positive and calbindin-negative principal cells (Figure 5C, *left* and *right* panels, respectively) in normal and Lis1 mutant littermates (5C vs E and 5D vs F, for PV and CB1R respectively). First, putative inhibitory boutons are automatically identified in the corresponding stain (Pv or CB1-R, top panels, blue staining). These putative pre-synaptically localized boutons are then filtered by proximity to a postsynaptic inhibitory synapse marker, gephyrin – yielding ‘true’ inhibitory puncta (synthetic spheres in bottom panels, gephyrin staining not shown). These puncta are then counted if they are within 0.2 μm or less of a principal cell soma – which are demarcated by the WFS1 antibody (green). Six calbindin positive and six calbindin negative principal cells in CA1 of mutants and non-mutant littermates are used for each image, yielding a single data point. The counts on calbindin-positive somas are divided by counts on calbindin-negative somas yielding a bias ratio (no. of Calb^+^ PCs / no. of Calb^-^ PCs). Numbers greater than one indicate a preference for calbindin-expressing principal cells.

PV-expressing basket cells preferentially innervated calbindin-negative principal cells in both mutant and non-mutant mice (Figure 5G; PV-calbindin preferences: 0.74 ± 0.05, 0.63 ± 0.11 for normal type and mutant respectively, p = 0.55, each point represents 12 cells from a slice, n = 3 pair of littermates from 3 litters). In experiments where the PV channel stain was replaced with a Cb1-R antibody, known to selectively stain presynaptic terminals of CCK-expressing basket cells, we noted a preferential innervation of calbindin-expressing post-synaptic targets in normal type that was absent from the Lis1 mutant mouse (Figure 5H; CB1-R-calbindin preferences: 1.32 ± 0.04, 1.02 ± 0.09 for normal type and mutant respectively, p = 0.02). Which suggested that at least from an immunohistological level, CCK-expressing basket targeting onto ectopic calbindin positive principal cells was disrupted.

### Monosynaptic CCK-mediated inhibition onto calbindin-positive principal cells is disrupted in CA1 of the Lis1 mutant

In order to better understand the role of CCK-expressing inhibitory cell networks in the face of pyramidal cell heterotopia and to further the observations shown in Figure 5 at a functional level, whole-cell recordings were made from principal cells in CA1 in the presence of excitatory synaptic transmission blockers (APV 50 uM and DNQX 10 uM). Monosynaptic inhibitory events were evoked using a stimulation electrode placed locally in the PCL of CA1, and omega-conotoxin (1 μM) was applied to selectively inhibit vesicle release from CCK-expressing interneurons (Figure 6) [37]. Example traces from four groups are shown in Figure 6C, from left to right, Wt complex, Wt simple, Lis1^+/−^ complex, Lis1^+/−^ simple. Baseline events are in black, and post wash-in data are in gray. In littermate controls, conotoxin reduced monosynaptically evoked IPSCs to 52.5 ± 3.9 % of baseline amplitudes in complex cells, while events in simple cells were reduced to 75.6 ± 8.3 % of baseline amplitudes, consistent with our observation that complex cells are preferentially targeted by CCK-containing interneurons (Figure 6D (*left*), p = 0.03, n = 8 Wt and 8 Lis1^+/−^ cells). In Lis1 mutant mice this differential CCK-containing inhibitory input was not detected, as conotoxin reduced eIPSCs to 48.2 ± 16.4 % of baseline and 60.2 ± 7.8 %, for complex and simple cell subtypes respectively (Figure 6D (*right*), p = 0.53 n = 13 and 5).

**Fig 6.**
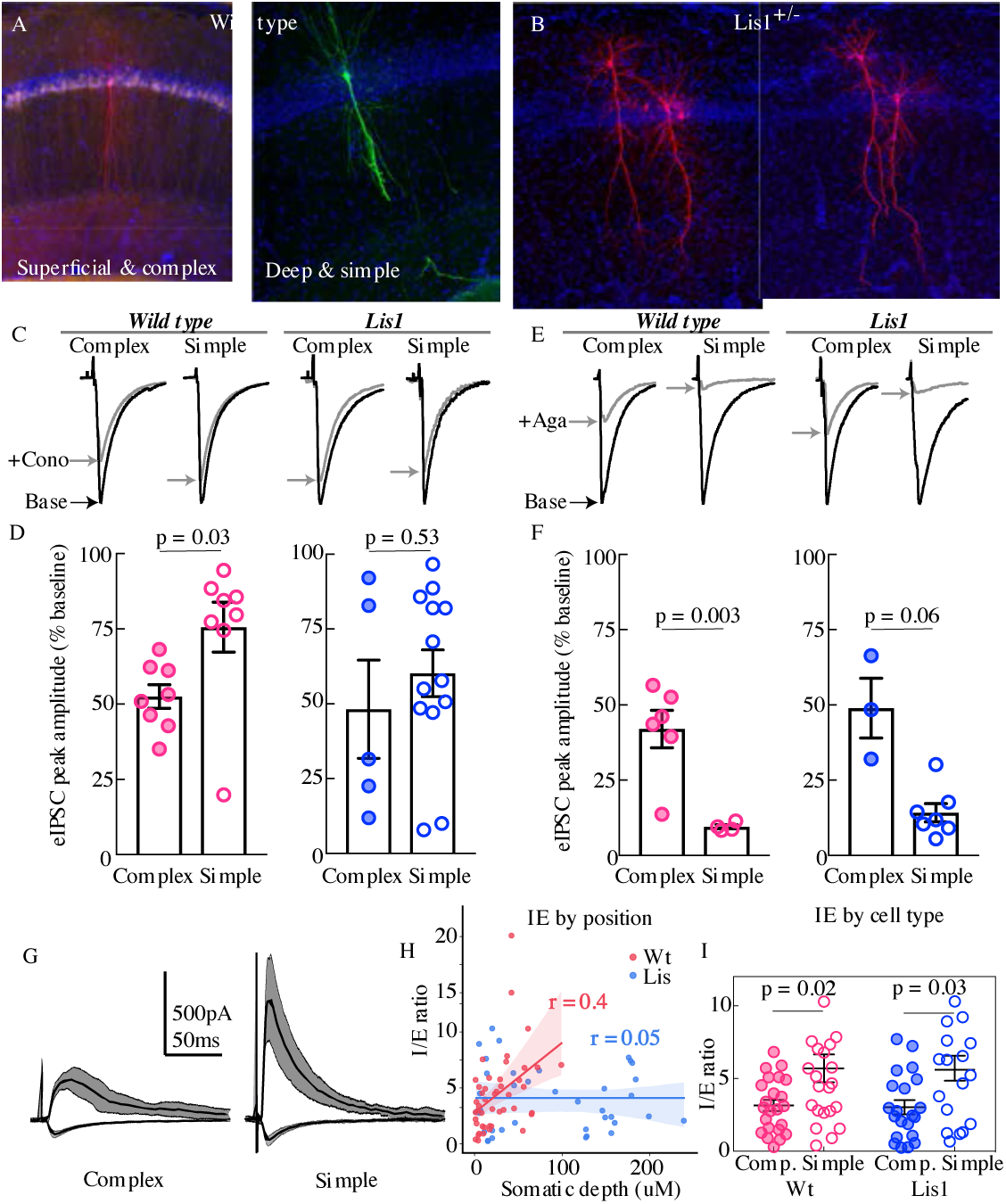
Physiological assays of network function within CA1. (A & B) Cell recoveries from normal type and Lis1 mutant experiments. (C) Normalized example traces from pre- and post-wash in (dashed) of omega-conotoxin (1 μM), *from left to right*, a normal-type complex and simple recordings, followed by Lis1^+/−^ complex and simple examples. Stimulation for monosynaptic experiments was delivered locally in the CA1 PCL. (D) Quantification of the percent reduction in the evoked IPSC 10-12 mins after drug application. Wt: complex 52.5 ± 3.9 %, simple 75.6 ± 8.3 %; Lis1^+/−^: complex 48.2 ± 16.4 %; simple 60.2 ± 7.8 %, n = 8, 8, 13, 5, respectively. (E) Example traces as in (C) but for omega-agatoxin experiments (250 nM). (F) As in (D) but for agatoxin. Wt: complex 42.0 ± 6.2 %, simple 9.5 ± 0.7 %; Lis1^+/−^: complex 48.9 ± 9.9 %; simple 14.2 ± 3.0 %, n = 4, 6, 7, 3, respectively. (G) Example traces for monosynaptic EPSCs (excitatory, inward current), and disynaptic feedforward IPSCs (inhibitory, outward current) evoked by stimulation of Schaffer collaterals, from a simple and complex recovered cell morphology in normal type. (H) IPSC amplitude / EPSC amplitude plotted by somatic PCL depth. (I) Same data as in (H) sorted by cell sub-type. Wt: complex 3.15 ± 0.39, simple 5.70 ± 0.95; Lis1^+/−^: complex 3.02 ± 0.49 %; simple 5.03 ± 0.76, n = 23, 23, 21, 17, respectively.

We next repeated this experiment using an antagonist known to inhibit release from parvalbumin-expressing interneurons, omega-agatoxin IVA (250nM). Example traces for the four subtypes before and after agatoxin application are shown in Figure 6E (wash-in data in gray). In control mice, agatoxin reduced monosynaptically evoked eIPSCs to 42.01 ± 6.2 % of baseline in complex cells, events in simple cells were reduced to 9.5 ± 0.7 % of baseline amplitudes, signifying that events in simple cells were more dependent on PV-expressing basket cell input (Figure 6F (*left*), p = 0.003, n = 4 complex and 6 simple cells). In Lis1 mutant mice agatoxin reduced eIPSCs to 48.9 ± 9.9 % of baseline and 14.2 ± 3 %, for complex and simple cell subtypes respectively (Figure 6F (*right*), n = 3 and 7, p = 0.06).

Having probed monosynaptic inhibitory circuitry onto putative calbindin-positive and - negative cells, we next examined feedforward disynaptic inhibition onto CA1 principal cells in normal and Lis1 mutant mice. Superficial cells have been previously shown to exhibit a comparatively higher level of excitatory drive during feedforward circuit activation (large EPSCs per unit of IPSC, [14]). Cells were voltage clamped at −70mV and +10mV to measure the Schaffer collateral-mediated monosynaptic excitatory and disynaptic inhibitory drive (Figure 6G). Excitatory transmission was subsequently blocked (APV 50 μM and DNQX 20 μM), to allow the subsequent isolation of the disynaptic feedforward inhibitory drive from the total inhibitory component. Inhibition:excitation (IE) ratios were positively correlated with somatic depth in the PCL for normal-type littermates, but not Lis1 mutants (Figure 6H; Wt r = 0.4, Lis r = 0.05). When recorded cells were sorted by complex and simple morphologies complex cells had lower IE ratios in both normal and Lis 1 mutant mice (Figure 6I, Wt complex 3.15 ± 0.39, simple 5.7 ± 0.95, p = 0.02 n = 23 complex and 23 simple; Lis1 complex 3.02 ± 0.49, simple 5.03 ± 0.76, p = 0.03 n = 21 complex and 17 simple cells). While their resulting ratios were predictive of sub-type, neither EPSC or IPSCs alone were significantly associated with depth or cell subtype (data not shown). EPSCs displayed depth correlations of r = 0.16 and r = 0.07 for normal-type and Lis1 experiments, respectively. Neither excitatory nor inhibitory events differed significantly between principal cell shapes. IPSCs had a somatic depth correlation value of 0.2 non-mutant littermates and 0.01 for mutants.

### Lis1^+/−^ mice display robust extracellular oscillations but are less synchronous across heterotopia

Using extracellular oscillations measured in vitro we next sought to assay alterations in network level function resulting from the cellular heterotopia present in our Lis1 mutants. Both normal-type and Lis1 mutant slices were capable of producing robust gamma oscillatory activity (ranging from 18-50Hz), in response to application of 20 uM carbachol (Figure 7) [38, 39, 40]. Slices from Lis1 mutants produced slightly higher frequency gamma oscillations than non-mutants (Wt 24.9 ± 1.7 Hz, Lis1^+/−^ 31 ± 1.1 Hz, p = 0.005 n = 20 and 14, respectively) (Figure 7B-D). Subsequent addition of the synthetic CB1R agonist, WIN-55,212-2 (WIN) (2 uM), did not alter the peak frequency of the oscillations in normal type nor mutant recordings (Figure 7D) but caused a significant decrease in peak power in normal type recordings (Figure 7E)., but not in Lis1^+/−^ mice suggesting that CCK-networks in mutants are less involved in gamma oscillation generation than in normal-type littermates (Wt +WIN 0.93 ± 0.03 vs CCh alone p = 0.03, Lis1^+/−^ +WIN 1.02 ± 0.04 vs CCh alone p = 0.69; n = 20 and 14 non-mutant and mutant respectively).

**Fig 7.**
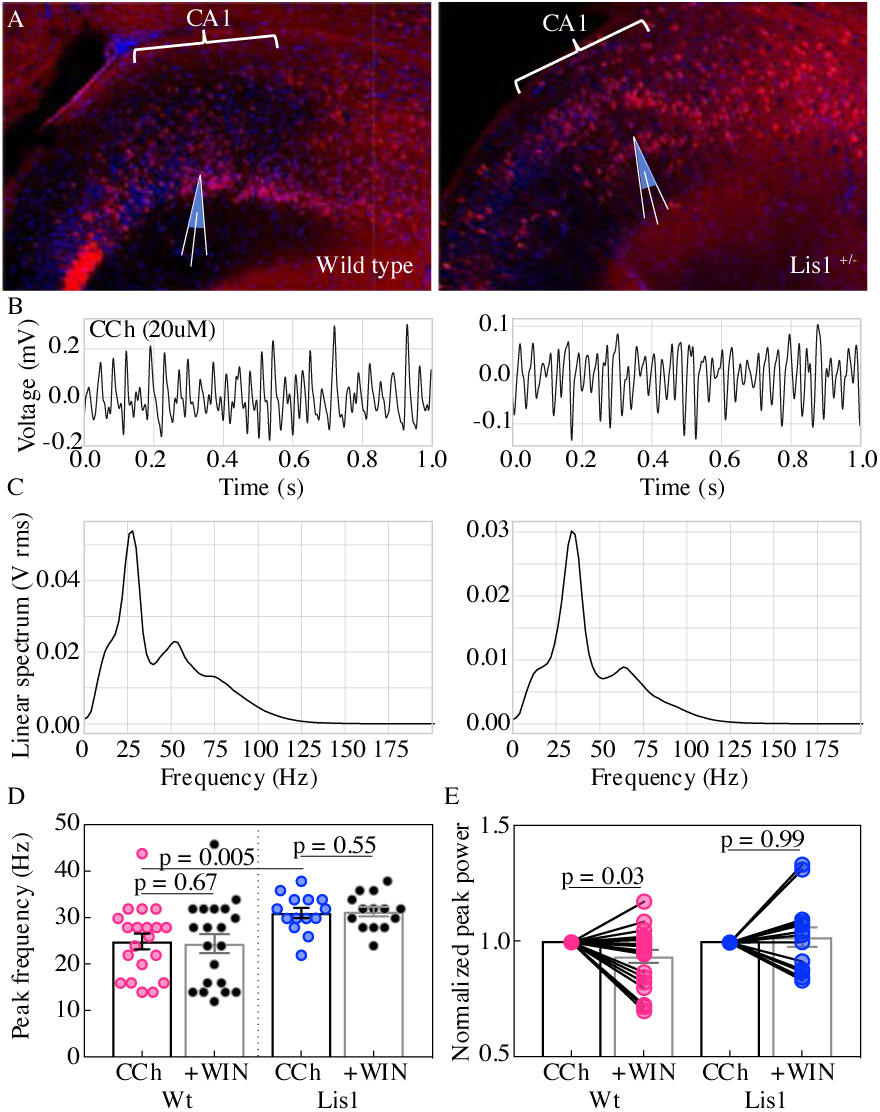
Lis1^+/−^ mice display robust carbachol induced oscillations. (A) Normal type (*left*) and mutant (*right*) images from ventral hippocampus in Calbindin-cre:Ai14 mice. Note the second layer of deeply positioned calbindin expressing principal cells in the Lis1 mutant. (B) One second of data during carbachol induced activity from radiatum side electrodes in normal type and mutant recordings, respectively. (C) Power spectra computed for each of the above example recordings. (D) Summary peak frequency data for non-mutant and mutant experiments, in carbochol alone, and with addition of WIN-55 (Cb1-R agonist, 2 um). Wt CCh 24.88 ± 1.7 Hz, +WIN 24.4 ± 2 Hz, Lis1^+/−^ CCh 31 ± 1.1 Hz, +WIN 31.3 ± 1 Hz. (E) Summary data as in (D) but for normalized Vrms power at the peak frequency. Wt +WIN 0.93 ± 0.03 vs CCh alone p = 0.03, Lis1^+/−^ +WIN 1.02 ± 0.04 vs CCh alone p 0.69; n = 20 and 14 non-mutant and mutant respectively. Pre-vs-post wash p values represent paired t tests.

In an additional series of experiments, a second electrode was placed in the same radial axis as the first approximately 150 um deeper, so that in normal type slices one electrode targeted the radiatum side of the PCL while the other targeted the oriens side (Figure 8A). In the Lis1 mouse slices electrodes were placed in different heterotopic bands but still in the same radial axis. This allowed for analysis of the correlation and synchronicity of oscillations across the normal and heterotopic layers of CA1 (Figure 8). Examples of simultaneous one second recordings are shown for the oriens (*top*) and radiatum (*bottom*) side electrodes in Figure 8B (Wt on *left*, Lis1^+/−^ on *right*). Dashed vertical lines show peak alignment for each example. Associated cross-correlation plots between these electrodes are displayed in Figure 8C (Wt *left,* Lis1 *right*); note the +0.7 ms peak in offset in the wild-type experiment, and −2.7 ms peak offset in the Lis1 example. Normal-type and Lis1 mutant slices were capable of producing correlated oscillatory activity (Figure 8D; Wt 394.6 ± 80.0, Lis1^+/−^ 394.2 ± 60.8, p = 0.99 n = 20 and 14). However, examining the time-shifts obtained from cross correlation analysis (how far one signal is peak shifted from another in time) we noted that Lis1^+/−^ mice displayed significantly less temporally correlated oscillations between the two electrodes (Figure 8E; Wt: +1.01 ± 0.8 ms, Lis1^+/−^: −1.8 ± 0.79, p = 0.02 n = 20 and 14) suggesting that while both heterotopic bands participate in the ongoing oscillation, their separation in anatomical space or deficits in basket cell network connectivity erodes the correlated activity between the bands. Application of the CB1R agonist WIN-55 produced modest decreases in non-mutant cross-correlation values but not in the Lis1 mutants (Wt: + WIN 333.9 ± 71.9, vs baseline p = 0.04 n = 20; Lis1^+/−^: + WIN 427.2 ± 84.13, vs baseline p = 0.43 n = 14) suggesting a diminished role for CCK-containing interneuron networks in the Lis1 mouse. WIN 55 application did not have a significant impact on the time-shift between deep and superficial channels in either genetic background (Wt: + WIN 0.68 ± 0.52 ms, vs baseline p = 0.62, Lis1^+/−^: + WIN −0.41 ± 1.23 ms, vs baseline p = 0.21).

**Fig 8.**
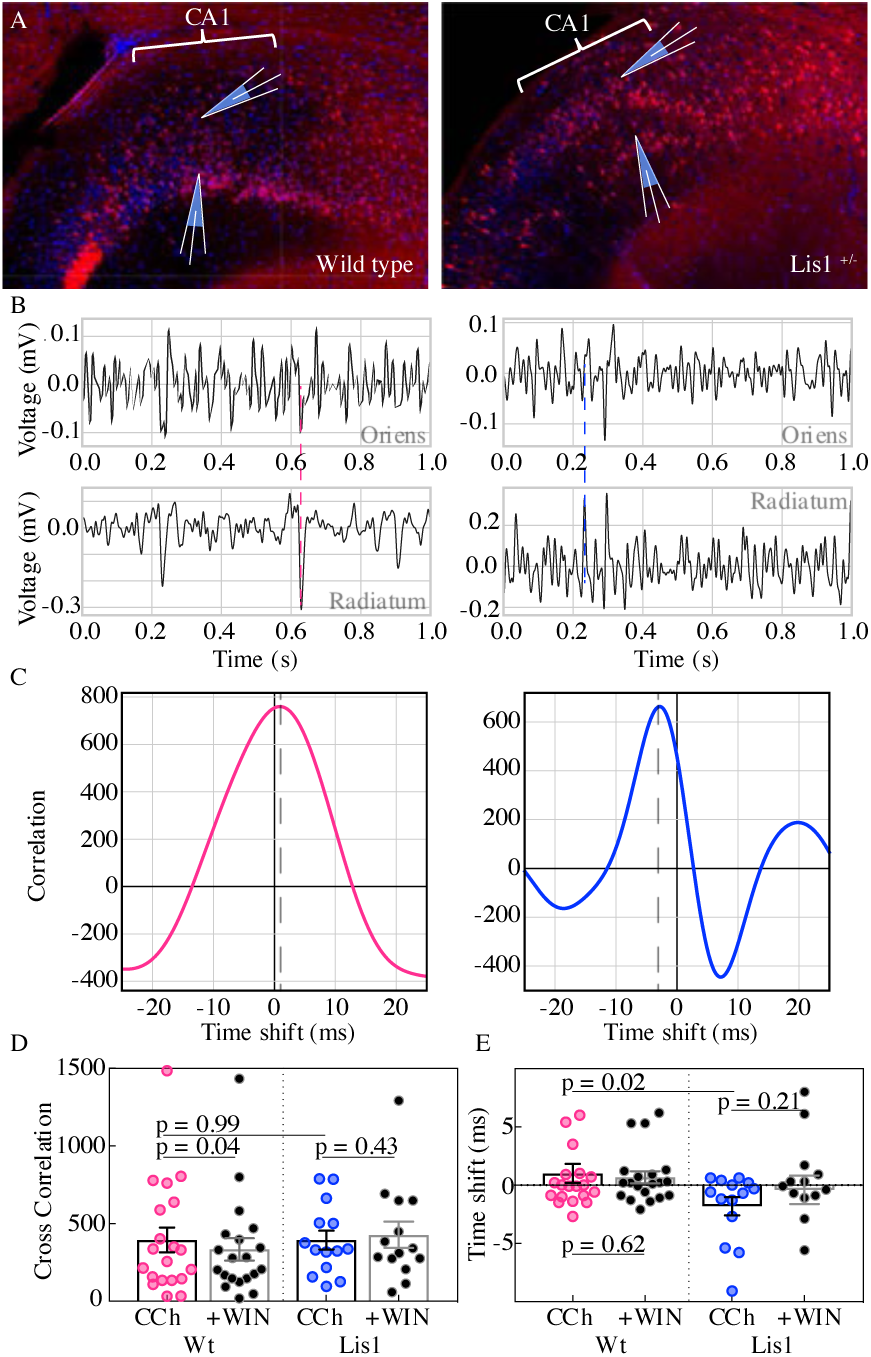
Carbachol oscillations in Lis1 mutants are less synchronous across CA1 heterotopias. (A) Normal type (*left*) and mutant (*right*) images from ventral hippocampus showing the positioning of dual electrode recordings, one from the s. radiatum and a second s. oriens side electrode in the same radial plane. (B) One second of simultaneous recordings from the deep (top) and superficial (bottom) electrodes, for non-mutant (*left*) and mutant (*right*) example experiments. Dashed lines highlight peak alignment between electrodes – note the blue line intersecting near a trough in the top trace, and a peak in the bottom. (C) Cross correlation plots for the example experiments shown in (B). Correlation values are arbitrary units. (D) Summary data for non-mutant and Lis1^+/−^ experiments in carbachol and after WIN-55 wash-in. Wt CCh 394.6 ± 80, +WIN 333.9 ± 72, Lis1^+/−^ CCh 394.2 ± 60.8, +WIN 427.2 ± 84.1. (E) Summary for the millisecond timing of peak correlation shifts shown in (D). Wt CCh 1 ± 0.8 ms, +WIN 0.68 ± 0.5 ms, Lis1^+/−^ CCh −1.8 ± 0.8 ms, +WIN −0.4 ± 1.23 ms; n = 20 and 14 non-mutant and mutant respectively. Pre-vs-post wash p values represent paired t tests.

## Discussion

Cellular heterotopias arising from various genetic and environmental factors carry with them a poor prognosis for the affected individual, including severe mental disability, increased seizure risk, and shortened life span [41]. The degree to which these effects are a direct result of the heterotopia itself (a lack of layers) or related to the role of the mutated genes in other processes remains unclear. That is to say, it is unknown to what extent any of the disease phenotypes associated with Lissencephaly are the result of disrupted layering and cellular misposition during embryonic development.

In the present work we first investigate the heterotopic banding observed in area CA1 of the Lis1 mutant mouse in order to determine if there is a pattern to the splitting of these excitatory cell populations. To this end we demonstrate that calbindin expressing principal cells are preferentially affected by cellular heterotopia in CA1, where they are proportionately relegated to the deeper cellular layer – opposite of their normal superficial positioning in the PCL (Figure 1). After confirming that these cells are the same embryonically derived population (Figure 2), namely late-born calbindin expressing, we asked to what degree their intrinsic development reflected the differences between calbindin-positive and calbindin-negative PC subtypes in normal type animals, and if relative differences between the two population were preserved (Figure 3 and 4). While there was an effect of stunted arborization in comparison to normal type calbindin cells, Lis1 calbindin cells retained their complex morphology relative to with-in animal non-calbindin expressing principal cells. Intrinsic physiological properties appear more disrupted in Lis1 calbindin expressing principal cells, however several properties showed greater differences or trended toward significant differences when separated by putative calbindin expression, as opposed to somatic positioning – suggesting again that subtype was a stronger influence than layering in the determination of these properties. It is unclear if the intrinsic physiological differences between calbindin positive PCs in normal and Lis1 mutants reflected other roles of the Lis1 protein directly, compensatory changes of ectopic cells, or are the result of cellular development in an ectopic position – though the first two seem more likely given findings from other mis-lamination models [26, 19, 21], though insufficient circuit integration and activity is known to alter interneuron development in cortex [42].

We next turned our attention to the integration of these ectopic calbindin expressing principal cells into the CA1 basket cell network. Staining experiments suggest that CCK expressing basket cell synapses were specifically altered to a greater extent than PV networks onto calbindin principal cell targets (Figure 5). This finding was confirmed by monosynaptic inhibition experiments that showed reduced sensitivity of ectopic calbindin expressing principal cells to a CCK cell antagonist, omega-conotoxin (Figure 6, left). Conversely, PV cell networks seemed substantially more resilient, which is not so surprising given that these cells occupy deeper positions within CA1, and their preferred synaptic targets are not substantially mispositioned under the cellular heterotopia present in Lis1 (Figure 5A and B) [11].

Disynaptic inhibition experiments support the notion of PV networks being more robust under cellular heterotopia (Fig 6, *right*). Feed-forward inhibition is much stronger onto PV baskets than their CCK expressing counterparts, making this largely a test of PV network connectivity [43]. Additionally, depolarization to +10 mV (as done in the experiment) drives depolarization-induced suppression of inhibition in CCK-basket cells, largely removing them from this assay [44, 45, 46]. In sorting these experiments by principal cell sub-type, we observed that ectopic calbindin expressing principal cells retained their relatively high excitability (low I/E ratios), suggesting that parvalbumin cells did not start to inappropriately target deeply positioned, ectopic calbindin PCs. Groups working in a related model of cellular heterotopia, the Reeler mouse which has severely disorganized cortical and hippocampal principal cell layering, previously reported that excitatory and inhibitory cells are produced in approximately the correct proportions, that ectopic cells retain expression of their correct markers, morphology of cell types is generally conserved, and their intrinsic physiological properties are largely unperturbed on a network level [22, 23, 47, 21]. Despite differing genetic causes, the present study supports these findings that brain development is surprisingly robust despite mis-lamination. An interesting caveat, however, is that in the present work, and studies of other cellular heterotopias, morphological development and orientation of principal cell dendrites appear stunted and meandering (Figure 3) [27, 34]. In the Reeler mouse synaptic network development was also remarkably intact, as thalamocortical and intracortical connectivity, cellular tuning properties to stimuli, and even animal behavior seem only minorly altered if at all [26, 19, 20,21]. From a broad perspective, this is in agreement with the present work in the Lis1 hippocampus, as feed-forward properties onto PC subtypes retain their relative excitabilities, and Lis1 slices retain their ability to generate gamma oscillations (Figure 7).

Interestingly, we observed higher peak oscillation frequency in Lis1 mutant experiments than normal type (Figure 7D). One possible interpretation of this result is that CCK-expressing interneuron networks tends to prefer lower frequency gamma, and when disrupted in Lis1 mutants networks become more dependent on alternative faster oscillation mechanisms such as greater reliance on parvalbumin cell networks. These results may reflect biological differences in hyperexcitability that predispose these mice and human patients to seizures. In the power domain, measurements are sensitive to differences in electrode placement between experiments, as this cannot be ruled out particularly as the cell layer positioning is unruly in Lis1^+/−^ mice, power data from these recordings was normalized and only compared within experiment to wash-in values (Figure 7E). Non-mutant slices showed power decreases in the presence of the cannabinoid receptor agonist WIN-55, while Lis1 mutant slices were non-responsive to this compound. These data add to our immunohistochemistry and monosynaptic physiology experiments in suggesting deficits in the CCK-basket cell networks of CA1 under heterotopia as Lis1 slices are largely not affected by WIN-55 application.

Comparing recordings from two electrodes in Figure 8 revealed that cross correlation values were relatively similar between normal type and mutant mice, but time-shifts or synchronicity between channels were significantly different (Figure 8E). It seems likely that timing differences in gamma-oscillations arise from the physical separation of current sinks and sources under Lis1^+/−^ heterotopia, and not as a result of the CCK-innervation deficit described above, as these measures were largely unchanged by WIN-55 application in normal-type mice, however that possibility cannot be ruled out [48, 49]. It is worth noting that the time-shifts under baseline conditions in the mutants are opposite in direction than that of non-mutants. In that respect, they roughly mirror the physical inversion of PCL lamina under Lis1^+/−^ cellular heterotopia.

Collectively, these findings bolster the notion that layers are in large part an epiphenomenon of neurogenesis, as has been hypothesized previously. Importantly, layer terminology has a correlated genetic component in normal type mice as it is likely to capture a related embryonic pool of neurons. Therefore, when traditional studies refer to cellular layer, they are using it as a proxy for cellular genetic subtype, which is no longer the case in heterotopias [26, 27, 24]. In agreement with this line of reasoning, decades of work on synapse development are increasingly bolstering the “hand-shake hypothesis” – where in molecular cues present on the surface of both putative synaptic partners confirm or reject synapse formation to aid in the establishment of appropriate and canonical circuitry over several scales of axon pathfinding [50, 51, 28, 29]. The degree to which these genetic network wiring mechanisms are modified in activity-dependent fashion afterword remains an area of active study [42, 52, 30]. Importantly, the present study does identify a crucial network motif, CCK targeting of calbindin positive principal cells, that is disrupted in ectopic calbindin PCs in the Lis1^+/−^ mouse. Further work will be needed to determine if this is a genetically specified connection preference for calbindin expressing principal cells, and why it might exhibit positional dependence.

It might not be so surprising to find specific defects in CCK-expressing synaptic connections as opposed to PV circuitry. CCK and PV expressing interneurons arise from different progenitor pools, in the caudal ganglionic eminence (CGE) and medial ganglionic eminence (MGE), respectively [53, 54]. Additionally, CGE interneurons are developmentally lagged relative to MGE pools, as MGE cells are born first [55]. Notably, later born basket cell populations (CCK basket cells), appear to be biased towards innervation of late born principal cell populations (superficial, calbindin expressing) in non-mutant animals. In fact, prior work has demonstrated that basket CGE derived populations wait until the first post-natal week to form synapses on principal cell somas in the PCL [56]. This network motif may represent a lopsided obstacle in the establishment of CA1 circuitry, as few if any of their putative synaptic targets remain on the radiatum adjacent side of the PCL under this form of cellular heterotopia [57]. As CCK cell somas reside largely on the border between the PCL and the radiatum, in the Lis1 hippocampus these basket cells are tasked with sending axons through the denser superficial PCL and passing through the inter-PCL space before finding their appropriate synaptic targets in the deeper heterotopic band. It remains to be seen whether this CCK specific defect is generalized to area CA1 in other cellular heterotopias, or Lis1 specific, but it may suggest natural limits to the handshake hypothesis – after all if you are never introduced, you cannot shake hands.

## Acknowledgements

J. D’Amour is supported by the National Institutes of General Medical Sciences (NIGMS) Postdoctoral Research Associate (PRAT) fellowship, award number Fi2 GM123992. C. McBain is supported by the Eunice Kennedy Shriver National Institute of Child Health and Human Development. The authors acknowledge with gratitude S. Hunt and D. Abebe for their assistance in tissue processing and animal management, M. Craig for help with analysis of oscillation experiments and K. Pelkey, R. Chittajallu, T. Petros, S. Lee, W. Lu. for feedback, comments, suggestions, and discussions during lab meetings. Finally, we are thankful to Dr. Wynshaw-Boris and his lab for providing the heterozygous floxed Lis1 mouse used to rederive the full het animal used here.

## Statement of competing interests

The authors declare no competing interests with this manuscript.

